# Comprehensive *Invitro* and *Insilico* Analysis of Secondary Metabolites from *Datura metel*: Promising Anti-Alzheimer’s Therapeutics

**DOI:** 10.1101/2024.03.21.586063

**Authors:** Meenakshi Sharma, Mukul Jain, Nil Patil, Abhishek Barnwal, Sumeet Tayade, Anil Kumar Delta, Chittaranjan Kole, Prashant Kaushik

**Author notes:** **Correspondence:** Mukul Jain, Prashant Kaushik (,). Equal author (Meenakshi Sharma, Mukul Jain).

## Abstract

This research investigates secondary metabolites from *Datura metel* as potential anti-Alzheimer’s therapies. In vitro techniques isolated extracts for AD pathology targeting, with in silico analysis identifying gene targets for prevention. Apigenin, Luteolin, and Withanolide A were studied, each with 300 potential gene targets and core gene counts of 54, 52, and 58 respectively. Lipinski’s Rule assessed their pharmacological properties, showing good absorption but limited blood-brain barrier penetration. Protein interaction mapping revealed shared targets among the compounds. GO enrichment and KEGG pathway analysis highlighted their impact on biological processes and pathways, suggesting their anti-Alzheimer’s potential. Luteolin notably reduced Aβ1–42 levels by up to 35.2% (*p<0.05) in SH-SY5Y cells, positioning it and Withanolide A as promising multi-functional Alzheimer’s medications. These findings underscore the significance of Datura phytochemicals in AD prevention and treatment.

## Introduction

Alzheimer’s disease (AD) stands as the most common form of dementia in the elderly, marked by a gradual deterioration in cognitive function and memory. Some of the leading causes of AD are the formation of extracellular amyloid-β plaques, the accumulation of abnormally phosphorylated tau within neurons, synaptic dysfunction, and neuronal cell death as typical events [1]. As of now therapies for AD predominantly involve the use of cholinesterase inhibitors, such as tacrine, donepezil, Rivastigmine, and galantamine, along with the non-competitive N-methyl-D-aspartic acid (NMDA) receptor inhibitor, memantine [2]. In the pursuit of countering Alzheimer’s disease-related factors, natural bioactive products emerge as crucial players. Earth harbours around 500,000 plant species, with roughly 120,000 of them holding promising potential to produce biologically active compounds. These compounds are pivotal in potentially shaping treatments for a wide range of diseases [3]. *Datura*, a genus within the *Solanaceae* family native to the Americas, encompasses 14 distinct species. Their natural habitat spans from the southwestern United States to the northern regions of Central America. [4] *Datura*, a plant in the Solanaceae family, derived its name from Sanskrit word ‘Dhutra’(divine inebriation) and has a rich history in healing. [5]. The *Datura* species are divided into three sections: *Datura* (3 species), *Dutra* (10 species) and *Ceratocaulis*, consisting of one species, respectively. [4] out of which only nine species are widely acknowledged: *D. ceratocaula Ortega, D. discolor Bernh, D. ferox L., D. innoxia Mil, D. kymatocarpa Barclay, D. leichhardtii Benth, D. metel L., D. quercifolia Kunth, D. stramonium L.,* and *D. wrightii Regel, D. metel* is found in Asia and Africa. Its roots contain atropine, while the aerial parts are abundant in scopolamine. [6]. The *D. metel* plant bears lengthy white flowers with a fragrant purple hue that span up to 6 inches, alongside broad, green, bread-shaped leaves measuring between 10 to 20 cm in length and 5 to 18 cm in width. Its fruits resemble spiny capsules, typically measuring 4-10 cm in thickness [7]. *D. metel* possesses a bitter taste and is acknowledged for its qualities as an anesthetic, anti-asthmatic, antispasmodic, anti-tussive, hallucinogenic, and hypnotic plant. Its dried seeds are regarded as a stronger soporific compared to the leaves. [8]. *Datura metel L*. contains key compounds like withanolides, alkaloids, flavonoids, sesquiterpenoids, lignans, and phenolic acids. Withanolides offer various health benefits, including immune regulation, fighting cancer, neuroprotection, blood sugar control, and potentially aiding Alzheimer’s disease [9]. Apart from alkaloids*, Datura* also synthesizes notable metabolites like flavonoids, the main phenolic compounds in the phenylpropanoids class, comprise subgroups like flavones (e.g., apigenin, luteolin), flavonols (e.g., kaempferol, quercetin), flavanones (e.g., naringenin, eriodyctiol), flavanonols (e.g., dihydroquercetin, dihydromyricetin), and isoflavones (e.g., daidzein, genistein) **Table 1** [10]. Flavonoids may positively impact the brain by interacting with glial signalling and neuronal pathways. This interaction has the potential to boost neuronal regeneration, enhance existing functions, protect vulnerable neurons, and influence the cerebrovascular and peripheral systems [11]. In context to previous research and studies this compound tends to have a significant role against Alzheimer disease. Luteolin has shown strong neuroprotective effects in various studies, defending against cognitive decline in rat models due to chronic cerebral hypoperfusion and obesity-related issues. It also reduces Aβ generation in Swedish amyloid precursor protein overexpressing cells and effectively mitigates AD pathologies induced by traumatic brain injury in recent research [12]. Luteolin at 50 μM inhibits TNFα-induced inflammation in HepG2 cells via NF-κB and AP-1 pathways. At varying doses (3.13–50 μM), it reduces 6-hydroxy-dopamine toxicity in PC12 cells, showing neuroprotective potential against neurodegenerative diseases like Alzheimer’s disease [13]. Multiple research groups have documented apigenin’s anti-inflammatory actions across various human and animal cell lines and models. Specifically, in a double transgenic mouse model APP/PS1, apigenin displayed promising neuroprotective potential. It notably improved memory impairments associated with AD, reduced the build-up of Aβ plaques, and countered oxidative stress [14]. In a study exploring apigenin’s anti-inflammatory and neuroprotective properties, it was evidenced that apigenin, at tested levels, does not exhibit neurotoxicity and presents neuroprotective capabilities. It diminishes caspase 3 activation, augments neuronal viability, primarily by modulating the inflammatory responses of microglia and astrocytes [15]. In the exploration of the complex features of *D. metel L*. and its various compounds, High-Performance Liquid Chromatography (HPLC) stands out as a leading tool in the field of phytochemical analysis [16]. The criticism of natural source drug discovery led to the rise of in-silico-directed real-time screening as a solution. Literature suggests that traditional and virtual high throughput screenings produce comparable bioactive hits. This method provides an alternative to expensive and labour-intensive in-vitro procedures, preventing resources from being allocated to compounds predicted as ineffective in in-silico analysis [17]. Network pharmacology represents a relatively recent method for methodically and comprehensively exploring how drugs exert their mechanisms of action [18]. Network pharmacology is an integrative computational method that constructs a network linking “protein-compound/disease-gene” interactions to unveil the mechanisms driving the synergistic therapeutic effects of traditional medicines [19]. Additionally, it aims to uncover fresh drug candidates and targets and repurpose existing drug compounds for diverse therapeutic purposes through an impartial exploration of potential target areas [20]. Therefore, this investigation seeks to understand and quantify the bioactive compounds present in *Datura metel* through HPLC as it empowers researchers to delve into the intricate chemical compositions within plant species. This investigation not only uncovers the inherent variability of *D. metel* but also highlights its potential to make valuable contributions to the realms of medicine and beyond. Also, we investigated the therapeutic potential of *Datura metel* by in-silico analysis **Figure 1**.

**Figure 1.**
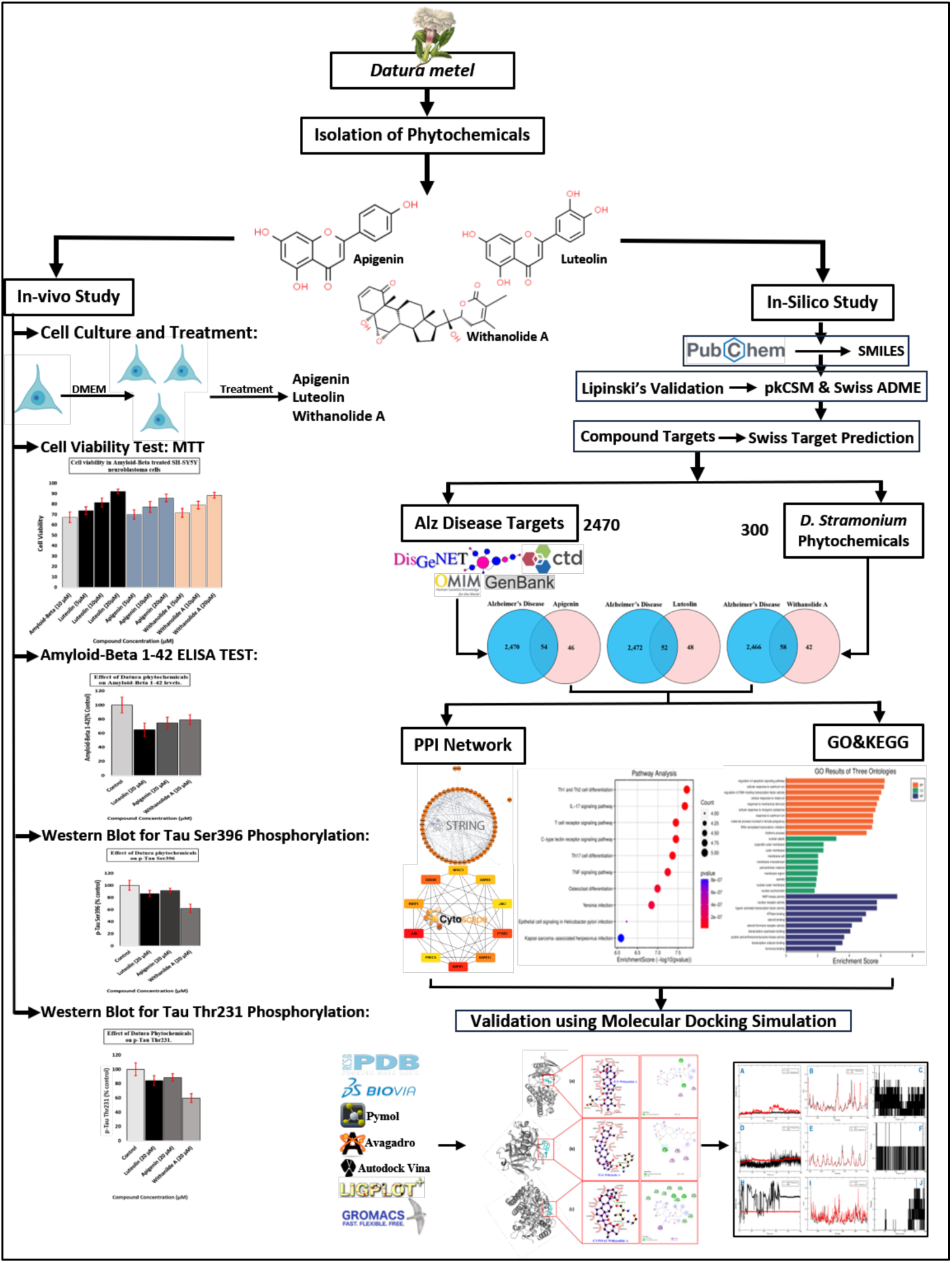
Systematic strategy workflow for the prediction of the preventive mechanisms of *Datura metel* phytochemicals in Alzheimer’s disease.

**Table 1.** Different bioactive compounds found in datura species and its invitro activity.

| Sr. | Species | Bioactive compounds | Activity | In-vitro | Ref |
| --- | --- | --- | --- | --- | --- |
| 1 | *Datura metel* | Daturalone | α-glucosidase and β-secretase inhibition. | Rat intestine powder | [7,21] |
|  |  | lignan-amides | reducie expression of iNOS and COX-2 | BV-2 microgilia cells |  |
| 2 | *Datura inoxia* | luteolin, apigenin, kaempferol, quercetin | inhibits TNFα-induced inflammation HepG2 cells | HepG2 cells | [22] |
| 3 | *Datura stramonium* | Atropine, scopolamine, amphetamine | Alters purinergic enzyme in neurotransmission system | Wistar strain albino rats | [23] |
|  |  | Leaf extract (DSL-EA) | Exhibits Anti-inflammatory Activity in CCL4-Induced Hepatic Injury Model | BALB/c mice | [24] |
| 4 | *Datura alba* | Magnesium oxide and zinc oxide | improve cognitive impairment and blood brain barrier leakage | BALB/C male mice | [25] |
| 5 | *Datura fastuosa* | Withanolides: A, B, C, E, baimantuoline A, B, C. | Anti-inflammatory | Rat paw edema. | [26] |
|  |  |  | Insecticidal effects: showed adverse effects on larval survival, weight and duration. | Helicoverpsa armigera |  |
| | | | Antimicrobial: effectively inhibited with bacterial concentration of 25μg/ml. | Escherichia coli. | |
| 6 | *Datura quercifolia* | Daturalactone, withahnicandrin, nicandrin, lycium, daturalactone 2 | Antioxidant: reduction of the DPPH radical and inhibition of lipid peroxidation. | Adult male Wistar rats | [4] |

## Results

### Concentration of compounds in seeds and leaves

The concentrations of six important biochemical compounds in seeds and leaves extracted with different solvents are presented in **Table S1**. The Scopolamine (mg/g) content in *D. metel* L. accessions ranged from 0.42 (D14) to 0.42 (D19) mg/g. Hyoscyamine (mg/g) content varied between 0.235 (D14) and 0.399 mg/g (D19). Atropine (mg/g) exhibited a range of 0.010 (D10) to 0.064 mg/g (D15). Withanolide A (mg/g) content spanned from 0.115 mg/g in D1 to 0.235 mg/g in D19 (Table 1). Similarly, the concentration of Withanolide B (mg/g) was lowest in D1 (0.095 mg/g) and highest in D19 (0.215 mg/g). Luteolin (μg/g) reached its peak at 12.529 μg/g in D8, while D19 had the highest concentration at 63.459 μg/g. Conversely, Apigenin (μg/g) was lowest in D25 (4.522 μg/g) and highest in D19 (43.449 μg/g). In conclusion, among the various Datura accessions, D19 exhibited the highest content followed by D15 **(Table S1)**.

### Analysis of variance (ANOVA)

The ANOVA outcomes for all assessed traits are detailed in **Table S2**. Evidently, variations attributed to accessions, environments, and the interaction between the two (A×E) were all notably significant (Table 3). This is in line with the anticipated complexity of phenotyping. The phenotypic coefficient of variation (PCV) estimates were slightly higher than the corresponding genotypic coefficient of variation (GCV), underscoring the environmental influence **(Table S2)**. Table 3 presents descriptive statistics for all accession traits. The coefficient of variation (CV) ranged from 7.380% for FL to 66.29% for Apigenin. Notably, five traits, mainly biochemical ones, exhibited CV values above 20%, while six traits had CVs between 10% and 20% **(Table S2).** Traits linked to biochemical tests demonstrated the highest CVs, whereas typically morphological traits (excluding Scopolamine) exhibited lower values. It’s noteworthy that traits with substantial quality significance, such as apigenin (66.29%), atropine (48.90%), luteolin (40.60%), withanolide B (23.81%), and withanolide A (20.85%), showed remarkable variability **(Table S2 & S3).**

### Data Processing of Core Gene Targets of Datura Phytochemicals and AD

The chemical structures of Datura phytochemicals i.e., “Apigenin, Luteolin, Withanolide A” were obtained from PubChem Database Online Server tool, and the SMILES files were taken for further inquiry. 300 potential Gene Targets were distinguished from each of the three potential phytochemicals in the table format from Swiss Target Prediction web tools. The Core Gene Targets for AD were found from CTD, DisGenNET, GenBank, OMIM database, with the relevance score ≥ 15 and score gda ≥ 0.2. At last a final number of 2524 Gene list was created by removing duplicates and extracted in .txt format using Venny2.1 (https://bioinfogp.cnb.csic.es/tools/venny/index.html) to visualize the inter-section result **(Table S4)**.

Now the common Gene Targets of AD and the Phytochemicals of Datura were overlapped using an Online tool Interactivenn (http://www.interactivenn.net/). Apigenin (54), Luteolin (52), and Withanolide A (58) in Venn Diagram **(Figure S1)**, implying that these genes were proven to be the core targets of Datura Phytochemicals for AD prevention and should be considered for subsequent PPI analysis.

### The Lipinski’s Rule for Absorption, Distribution, Metabolism, Excretion and Toxicity (ADMET)

The 2D structures of the probable phytochemicals from *Datura metel* were analysed for their pharmacological and molecular properties including molecular weight, oral bioavailability, drug-likeness, Caco-2 permeability, blood-brain barrier, fractional negative accessible surface area, log P, hydrogen bond donor and hydrogen bond acceptor, number of rotatable bonds, and topological polar surface area, which were all collected from the pkCSM server (https://biosig.lab.uq.edu.au/pkcsm/prediction) and Swiss ADME server (http://www.swissadme.ch/). Phytochemicals Apigenin, Luteolin, and Withanolide A were found to be poorly absorbed by the Blood Brain Barrier (BBB) **(Table S5)**. The intestinal absorption values of all these 3 Phytochemicals were greater than 30% making them a better absorbing compound from the intestine after oral administration, which indicates that all the natural compounds could be remarkably absorbed from the intestine of humans. Not all of the ligands were CYP1A2, CYP2C19, CYP2C9, CYP2D6 and CYP3A4 enzyme inhibitors. Only Apigenin can catalyse or inhibit CYP1A2, CYP2D6 and CYP3A4. CYP2D6 enzyme can catalyse basic compounds with protonated atoms 4-7 A such as several types of flavonoids and alkaloids. The CYP3A4 enzyme is a type of P450 that can catalyse most of the lipophilic active molecules. Cytochrome P450 enzymes in drug metabolism: regulation of gene expression, enzyme activities, and impact of genetic variation. Apigenin is at risk of narcosis or basic toxicity due to the presence of monocyclic ether reactive substances and not epoxides or peroxide. All of these functionalization reactions are biotransformation stages in phase 1 metabolism [Ideaconsult. 2011. Toxtree User Manual 5th Version. Sofia, Bulgaria.] Another toxicity parameter that needs to be known is the identification of certain substances or active groups that are label metabolized by the CYP3A4 enzyme using the SMARTCyp method. Prediction results showed that all the best compounds had certain active groups that were labile to metabolize by the CYP3A4 enzyme. There are several functional reactions identified in the three best compounds, ranging from aliphatic hydroxylation, aromatic hydroxylation, alcohol oxidation, O-dealkylation and epoxidation. Aliphatic hydroxylation is an oxidation reaction involving CN chain heteroatoms, while O-dealkylation is an oxidation reaction involving CO Chain heteroatoms. Epoxidation is a type of non-oxidizing reaction, that is, the hydrolysis type for the conversion of epoxides to side-by-side diols. Apigenin only undergoes aromatic hydroxylation reactions.

### Protein Interaction Network Map of Datura Phytochemicals

Protein-Protein Interaction forming a web like structure for various multifunctionality and dynamic roles of protein in various signalling pathways. So, to obtain the PPI networks of Apigenin, Luteolin, and Withanolide A with AD, the core targets of Apigenin (54), Luteolin (52) and Withanolide A (58) were extracted and paste into STRING database, and the visualization was performed using Cytoscape application (version 3.10.0). PPI in string network is a unique way of showing interaction with the desired protein of interest as the “Nodes” (Circles) in the Network represent various Targets molecules i.e., Genes and Protein to be very specific, and the edges (connecting lines) represent “Biological bridges linking proteins in a tapestry of interactions.” Supporting the understanding of the functional organization of the proteome and cellular process in AD progression. The PPI network of Apigenin-AD consisted of 54 nodes and 290 edges with an average node degree of 10.7. In this network of Luteolin-AD, 52 nodes, and 280 edges with an average node degree of 10.8 were included. In addition, the PPI network of Withanolide A-AD contained 58 nodes and 252 edges with an average degree of 8.69. The minimum interaction score was set to >0.4 and the Species was chosen as *Homo sapiens* **(Figure S2).** By using the CytoHubba plugin in Cytoscape (3.10.0) the nodes of each Datura phytochemicals were broken into the top 10 targets with the higher degree of connectivity and were selected as hub genes. Surprisingly the top 9 hub genes of Apigenin and Luteolin shared exactly nine genes MMP2, GSK3B, ESR1, KDR, AKT1, IGF1R, MMP9, EGFR, PTGS2 for AD prevention, and suggesting that they may share a common pathway for the treatment of AD. In case of Withanolide A MAPK1, MAPK8, JUN genes shows in the AD prevention network **(Figure S3)**. An UpsetR diagram was constructed to obtain an intersection between the top 10 hub genes of the three phytochemicals compound in AD. The results enlightened that the common targets of Apigenin and Luteolin are PARP1 and Luteolin-Withanolide A are PTGS2, GSK3B, and KDR suggesting that Luteolin-Withanolide A might have a similar mechanism of action, while Apigenin might exert somewhat different ones on AD prevention **(Table 2 & Figure S4)**.

**Table 2.** Molecular interaction between Datura phytochemicals and the common gene associated with it.

| Group | Interaction Number | Interaction Elements |
| --- | --- | --- |
| Apigenin | 6 | AKT1, EGFR, ESR1, MMP9, MMP2, IGF1R |
| Luteolin | 1 | MET |
| Withanolide A | 6 | JUN, MAPK1, MAPK14, NR3C1, MAPK8, JAK2 |
| Apigenin-Luteolin | 1 | PARP1 |
| Luteolin-Withanolide A | 3 | PTGS2, GSK3B, KDR |

### GO Enrichment and KEGG Pathway Analysis of Datura Phytochemicals

In this study, we explored the intricate mechanisms underlying the biological effects of Datura Phytochemicals of Apigenin, Luteolin, and Withanolide A through the analysis of Gene Ontology (GO) and Kyoto Encyclopedia of Genes and Genomes (KEGG) pathways. GO enrichment analysis provides valuable insights into the functional significance of genes associated with the target Phytochemicals. By categorizing gene functions into biological processes, cellular components, and molecular functions, we gained a comprehensive understanding of the potential impact of Phytochemicals on cellular activities. The top 10 enriched categories in Biological Process (BP), Cellular Component (CC), and Molecular Function (MF), in ascending order of p-values, are shown in the GO enrichment histogram **(Figure S5)**.

The core targets of Apigenin were involved in 924 BP terms, 36 CC terms, and 57 MF terms. In the BP analysis, the key target genes of Apigenin were mainly enriched in response to amyloid-beta, response to oxidative stress, cellular response to chemical stress, cellular response to reactive oxygen species, neuroinflammatory response, etc. The CC ontology identified significant enrichment for the platelet alpha granule lumen, membrane raft, membrane microdomain, membrane regions, etc. The MF category was mainly enriched with serine-type endopeptidase activity, serine-type peptidase activity, serine hydrolase activity, metalloendopeptidase activity, etc.

GO analysis of Luteolin-anti-AD showed that 875 BP terms, 38 CC terms, and 56 MF terms were enriched. Luteolin core targets were involved in response to amyloid-beta, response to oxidative stress, cellular response to chemical stress, cellular response to reactive oxygen species, neuroinflammatory response, etc. The core targets of Luteolin in CC were involved in platelet alpha granule lumen, platelet alpha granule, collagen-containing extracellular matrix, nuclear envelope lumen, astrocyte projection, etc. The MF analysis of Luteolin suggested serine-type endopeptidase activity, serine-type peptidase activity, serine hydrolase activity, metalloendopeptidase activity, endopeptidase activity, and others.

The core target of Withanolide A-anti-AD was found to be associated with 508 BP terms, 16 CC terms, and 46 MF terms. As shown in Figure, BP analysis results of Withanolide A were regulation of apoptotic signaling pathway, cellular response to cadmium ion, regulation of DNA-binding transcription factor activity, cellular response to metal ion, etc. The enriched CC ontologies of Withanolide A were dominated by the nuclear speck, organelle outer membrane, membrane raft, membrane microdomain, etc. In addition, factors such as MAP kinase activity, nuclear receptor activity, ligand-activated transcription factor activity, ATPase binding and others were included in the MF enrichment of Withanolide A.

Our investigation further extended to the exploration of KEGG pathways, elucidating the intricate networks of molecular interactions and signaling cascades influenced by our three Phytochemicals of Datura **(Table S6)**. KEGG pathways offer a systematic view of how these Phytochemicals may modulate key biological processes, providing a roadmap to comprehend the intricate interplay between genes and their functional implications. The top 10 pathways were chosen to build bubble maps with a threshold of p < 0.05. In total, 66, 64, and 83 significant pathways were exported for Apigenin, Luteolin, and Withanolide A respectively. Apigenin mainly acted on pathways as Proteoglycans in cancer, EGFR tyrosine kinase inhibitor resistance, and the IL-17 signaling pathways, while Luteolin principally affected the Proteoglycans in cancer, IL-17 signaling pathways, and Prostate cancer pathway. KEGG pathways for Withanolide A have principally enriched in Th1 and Th2 cell differentiation, IL-17 signaling pathways, and T cell receptor signaling pathway. In addition, IL-17 signalling pathway and TNF-signalling pathway were common pathways for these three phytochemicals **(Figure S5 & Figure 2)**. It can be said that the hub genes of *Datura metel* were closely related to top 10 pathways, demonstrating that these hub genes of our compounds might play a role in solving the unresolved AD prevention in the future.

**Figure 2.**
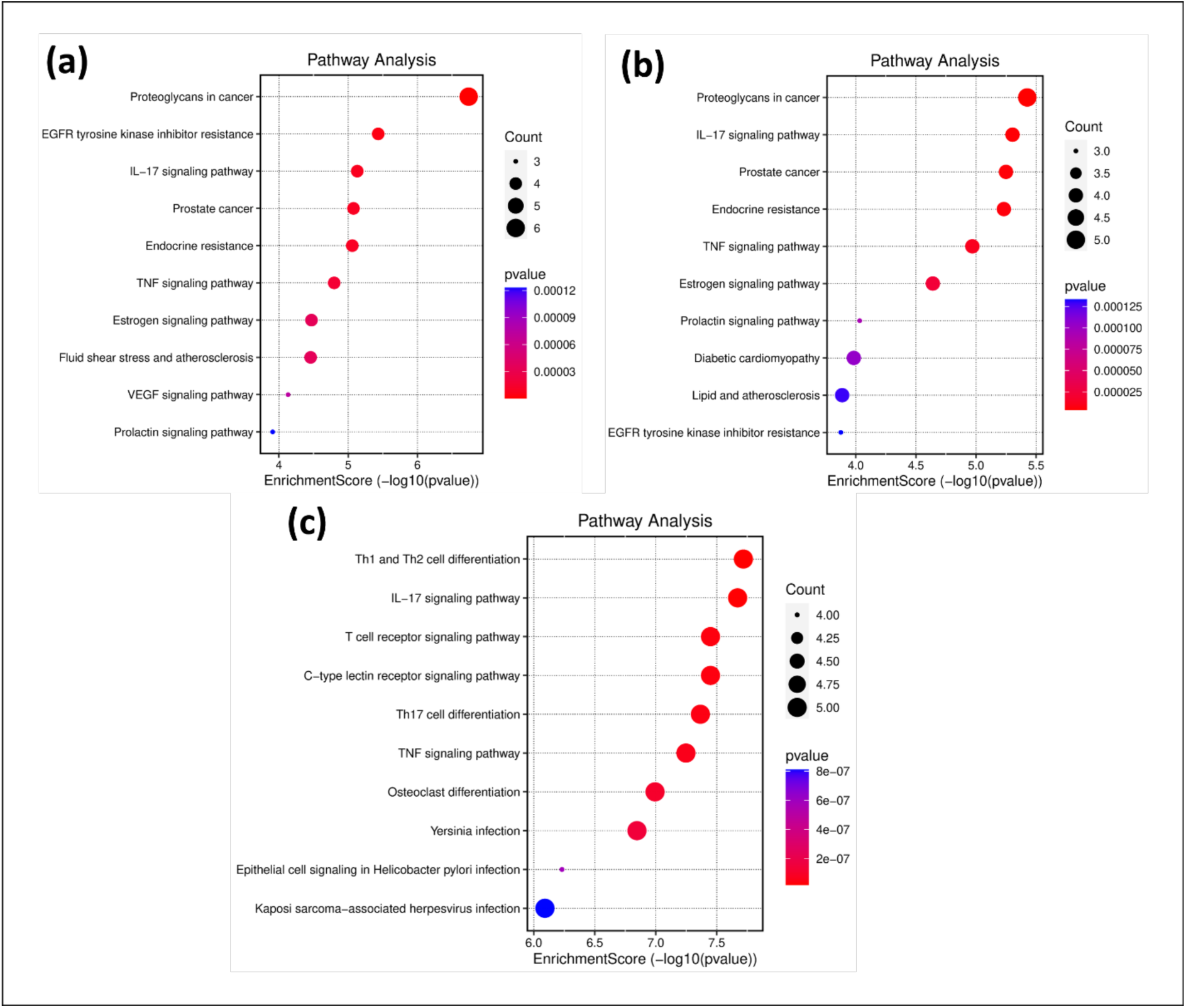
KEGG pathways enrichment analysis of **(a)** Apigenin, **(b)** Luteolin, and **(c)** Withanolide-A. The x-axis represents the gene ratio; the y-axis represents the enrichment pathway; the size of the dot represents the number of genes; the color of the dot represents the level of p-value.

### Molecular Docking Validation of Datura Phytochemicals

Since the active phytochemicals of *Datura metel* are three kinds i.e., Apigenin, Luteolin, and Withanolide A, and JUN, PLG, and CYP19A1 are the top three top-ranked hub genes to validate the network pharmacology results mainly screened by degree value among the intersecting targets and various literature articles and reviews, so that the molecular docking validation of the most important components and the most important targets has better credibility. By doing Molecular Docking of the top three-degree value of the core targets with the phytochemicals using AutoDock software the result shows significant strong binding affinity with one phytochemicals of Datura i.e., Withanolide-A with all three top-ranked hub genes as −9.1 with PLG (Plasminogen), −10.46 with CYP19A1 (Cytochrome P450 Family 19 Subfamily A Member 1), and −11 with JUN. The JUN-Withanolide-A complex formed hydrogen bonding with residues ALA74 and GLN75. The binding affinity of the PLG-Withanolide-A complex was attributed to a hydrogen bond with only one residue i.e., PHE748. CYP19A1 interacts with Withanolide-A by forming hydrogen bonding with GLY433 and LYS440. The results of Molecular docking and binding energies with interaction at the active site are presented in the **Figure 3 & Table S7** below.

**Figure 3.**
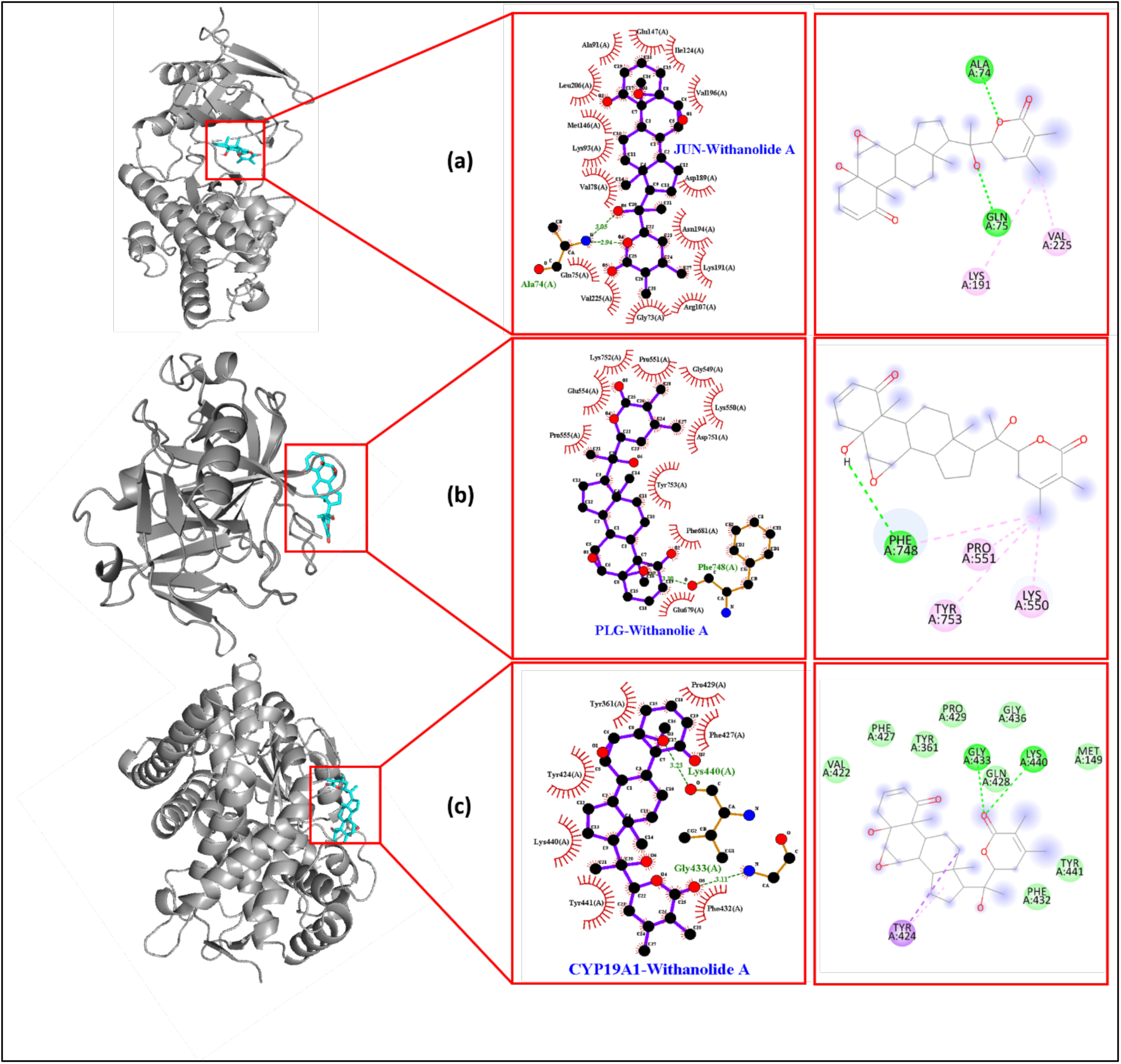
Molecular docking analysis of **(a)** Apigenin, **(b)** Luteolin, and **(c)** Withanolide-A. The interaction was seen via PyMOL and the 2-D Protein ligand interaction was visualize by Ligplot+ and Biovia Discovery Studio software.

### Molecular Dynamics validation of Datura Phytochemicals

#### Analysis of Molecular Dynamics Simulation

Molecular Dynamics (MD) modeling serves as a valuable tool for exploring protein-ligand complexes within a cellular environment, assessing the flexibility of the complex, and evaluating the stability of the protein following binding with a compound or ligand. The selection of the 2P33-WA (Complex 1), 3QEM-WA (Complex 2), and 3UIR-WA (Complex 3) complexes for a 100 ns Molecular Dynamics Simulation (MDS) was based on their robust binding interactions, alignment with drug-like characteristics, and favorable ADMET properties.

#### Root Mean Square Deviation (RMSD) Analysis

In the context of molecular dynamics simulations, the assessment of Root Mean Square Deviation (RMSD) serves as a valuable metric for gauging the stability and dynamic behavior of protein–ligand complexes. RMSD quantifies the deviation in the structure of a protein or protein–ligand complex from its initial conformation, offering insights into the complex’s stability during the simulation. In our study, we present an analysis of the RMSD behavior for 2P33, 3UIR, and 3EQM either in isolation or in complex with Withanolide-A (WA) during molecular dynamics simulations under physiological conditions **(Figure 4 A-C)**. Notably, the RMSD of 3QEM alone exhibited an abrupt increase in the absence of WA during the initial 2 ns, followed by a consistent behavior for the subsequent 98 ns. Conversely, the RMSD fluctuations of 3UIR and 3EQM alone remained within acceptable limits throughout the simulation. Furthermore, Complex 2 displayed fluctuations in the range of 15-17 nm until 65 ns, followed by a stable phase for the next 35 ns, which is deemed unacceptable. On the other hand, Complex 1 and Complex 3 exhibited slight fluctuations around 0.4 nm and 0.75 nm, respectively. These findings imply that the overall structures of the target enzymes (2P33 and 3UIR) did not undergo significant changes due to the binding of Withanolide-A. Consequently, the protein–ligand complexes remained stable over the course of the simulation.

**Figure 4:**
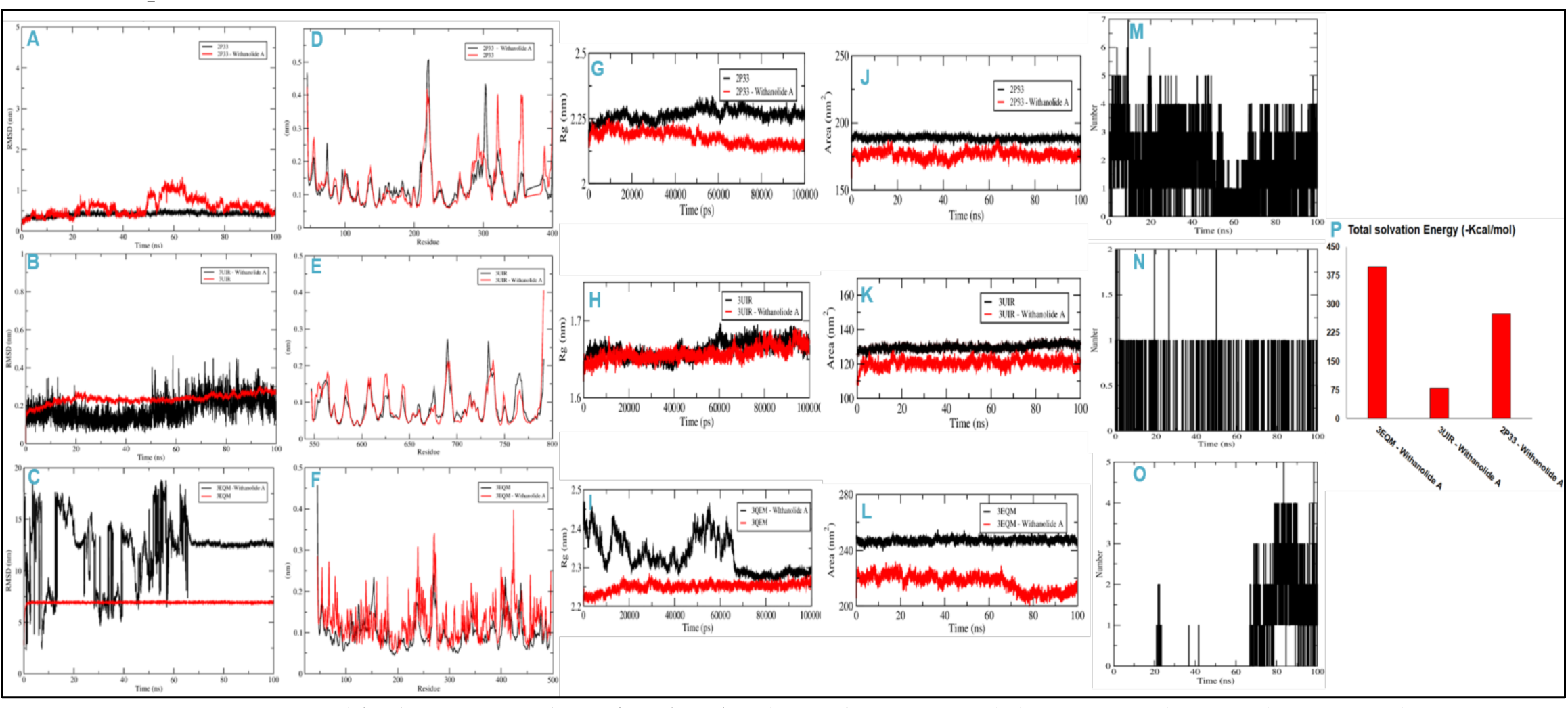
Graphical representation of molecular dynamics, RMSD (A), RMSF (D), Rg (G), SASA (J), Hbonds (M) of complexes 1; RMSD (B), RMSF (E), Rg (H), SASA (K), Hbonds (N) of complexes 2; RMSD (C), RMSF (F), Rg (I), SASA (L), Hbonds (O) of complexes 3; And Total solvent energy of all complexes (H).

#### Root Mean Square Fluctuation (RMSF) Analysis

During Molecular Dynamics Studies (MDS), assessing the Root Mean Square Fluctuation (RMSF) of proteins is crucial for understanding local conformational changes in side chains resulting from ligand binding. In our investigation, we tracked the RMSF of Withanolide-A bound to 2P33, 3UIR, and 3QEM **(Figure 4 D-F)**. Typically, higher fluctuations are observed in residues located at the N and C-terminal or loop regions. The average RMSF values for 2P33, 3UIR, and 3QEM in the presence of Withanolide-A were 0.5 nm, 0.4 nm, and 0.35 nm, respectively. These findings suggest that the RMSF of 2P33, 3UIR, and 3QEM did not exhibit significant deviations in the presence of Withanolide-A. Consequently, this indicates that the overall conformation of the target proteins was conserved, highlighting the stability of the protein-ligand complexes.

#### Analysis of Radius of Gyration (Rg) and Solvent Accessible Surface Area (SASA)

The examination of the radius of gyration (Rg) and solvent accessible surface area (SASA) of a ligand over simulation time provides insights into the ligand’s behavior within the enzyme’s binding pocket. Rg values articulate the Root Mean Square Deviation (RMSD) of an atom’s width from the common center of mass, offering a measure of whether the complex maintains its folded state during Molecular Dynamics (MD) simulation. In the presented study, the fluctuation in Rg values for Withanolide-A bound to proteins (2P33, 3UIR, and 3QEM) over simulation time is depicted in **(Figure 4 G-I)**. The results indicate that Rg values for different protein–ligand systems exhibited fluctuations within acceptable limits throughout the simulation. Notably, the average Rg values of 2P33, 3UIR, and 3QEM bound with Withanolide-A were reduce by approximately 50% compared to the proteins alone. This reduction in Rg values suggests a more compact structure of the protein–ligand complexes, emphasizing the stabilizing effect of Withanolide-A on the overall structure of the binding systems.

The solvent accessible surface area (SASA) is a metric that gauges a protein’s exposure to the solvent, providing insights into whether the protein maintains its native conformation upon ligand binding. In this investigation, we quantified the SASA of the target proteins 2P33, 3UIR, and 3QEM bound to Withanolide-A (WA) **(Figure 4 J-L)**. The analysis reveals slight variations in the SASA of the complexes, all within acceptable limits. The average SASA values for Complex 1, Complex 2, and Complex 3 were determined to be 185 nm, 127 nm, and 230 nm, respectively. These findings indicate that Withanolide-A consistently resides within the binding cavity of 2P33, 3UIR, and 3QEM, maintaining a stable conformation throughout the course of the simulation.

#### Contact between Withanolide A and Target Proteins

The confirmation of a stable protein and ligand complex formation was ascertained by quantifying the total number of contacts established between them over the course of the simulation **(Figure 4 M-O)**. It is evident that throughout the simulation, the total number of contacts in the form of hydrogen bonds between Withanolide-A and the proteins 2P33, 3UIR, and 3QEM exhibited variations, amounting to 7, 2, and 5 hydrogen bonds, respectively. These findings unequivocally validate the persistent occupancy of Withanolide-A within the binding pockets of the target proteins over the simulation duration.

Moreover, Complex 3 exhibited fluctuations in hydrogen bond formation until the 70 ns mark. Subsequently, a more stabilized pattern emerged, with the formation of 3-5 hydrogen bonds post the 70 ns timeframe. This nuanced observation underscores the dynamic nature of the hydrogen bonding interactions in Complex 3 during the latter part of the simulation. The establishment and maintenance of specific hydrogen bonds are indicative of the stability and specificity of the protein-ligand interactions. These interactions are crucial for the overall structural integrity and function of the protein-ligand complexes. The observed fluctuations and subsequent stabilization in hydrogen bond formation provide valuable insights into the temporal evolution of the molecular interactions governing the Withanolide-A and protein complexes. Moreover, the quantitative analysis of hydrogen bond dynamics contributes to our understanding of the intricate intermolecular forces at play, shedding light on the nature and strength of the interactions that define the stability of the protein-ligand complexes. This comprehensive investigation into the time-dependent hydrogen bond dynamics enriches our knowledge of the complex molecular mechanisms underlying the binding interactions between Withanolide-A and the target proteins 2P33, 3UIR, and 3QEM, elucidating the nuanced intricacies of their interaction profiles.

### Validation of the neuroprotective effects of *Datura metel* phytochemicals against Alzheimer induced SH-SY5Y neuroblastoma cells

In this experiment, the set of analyses examined the impact of Luteolin, Apigenin, and Withanolide-A on cell viability. The result shown in Figure 5A, indicates that in SH-SY5Y neuroblastoma cells, Luteolin at 20 µM increase the cell viability by 24.9% (*p<0.05). At 10µM, Luteolin increase cell viability only by 6.4% (*p<0.05). Apigenin at 20 µM increase cell viability by 18.5% (*p<0.05), while Withanolide-A at 20 µM increase cell viability by 21.2% (*p<0.05). Since all three compounds (Luteolin, Apigenin, and Withanolide-A) shows statistically significant increase in cell viability by using MTT assay, but the most significant cell viability was achieved by Luteolin phytochemicals of *Datura metel* plant **(Figure 5 A)**.

**Figure 5.**
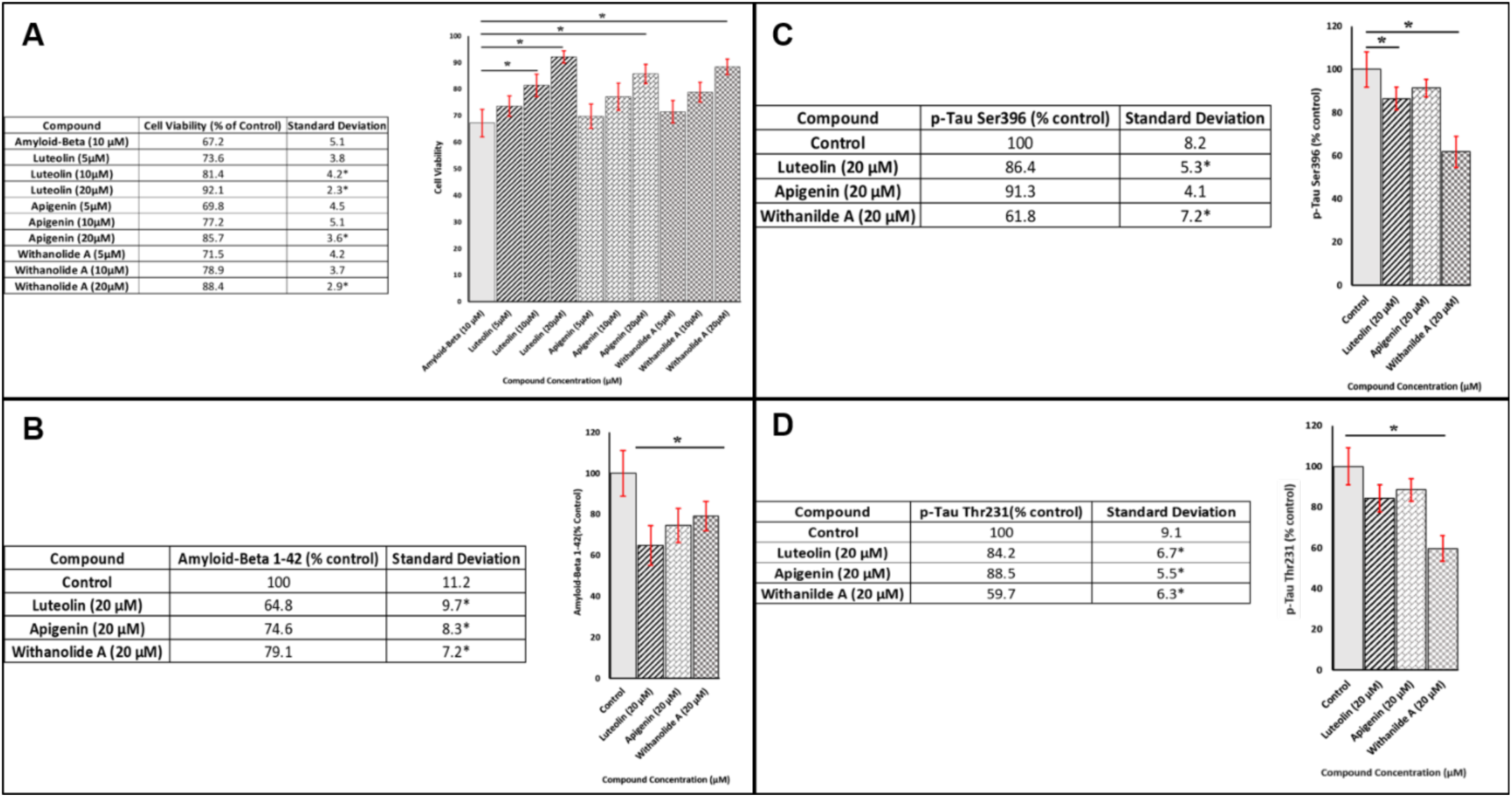
(A) Effect of Datura phytochemicals on cell viability in amyloid-beta treated SH-SY5Y neuroblastoma cells. (B) Effect of Datura phytochemicals on Amyloid-Beta 1-42 Levels. (C) Effects of Datura phytochemicals on Tau Phosphorylation, Western Blot result of p-Tau Ser 396 (D) Effects of Datura phytochemicals on Tau Phosphorylation, Western Blot result of p-Tau Thr 231.

The neuroprotective effects of (Luteolin, Apigenin, and Withanolide-A) in the SH-SY5Y cells were evaluated using ELISA technique for the estimation of Aβ_1–42_-induced cells in Figure 5B. A 20 µM of each three phytochemicals of *Datura metel* i.e., Luteolin, Apigenin, and Withanolide-A was used on the initial SH-SY5Y neuroblastoma cells. At these concentration Luteolin shows the high potency of reducing Aβ_1–42_ up to 35.2% (*p<0.05). At the concentration of 20 µM Apigenin and Withanolide-A when compared to the control doesn’t exhibit significant cytotoxicity in the SH-SY5Y cells (*p<0.05). To assess the neuroprotective effects of Luteolin, Apigenin, and Withanolide-A, the neuroblastoma cells were pre-treated with Luteolin, Apigenin, and Withanolide-A for 2h and incubated for 24h with 1 µM of Aβ_1–42._ Pre-treatment of the cells with Luteolin (20 µM) significantly protected (p<0.05), the SH-SY5Y cells from the Aβ_1–42_ when compared to the cells exposed only with the Aβ_1–42_ **(Figure 5 B).**

### p-Tau Ser 396

Previous In-silico study suggested a role of IL-17/TNF axis in tau pathology and AD development. The use of *Datura metel* phytochemicals Apigenin, Luteolin, and Withanolide-A, shows the significant decrease in the p-tau (Ser396) to tau, while the level of β-actin remained the same across all lanes. Treatment of tau phosphorylated SH-SY5Y neuroblastoma cells at the site of Ser396 shows that Withanolide-A reduces the tau hyperphosphorylation at Ser 396 with 38.2% significantly (*p<0.05). The other compound i.e., Luteolin and Apigenin has overall reduction of 13.6% and 8.7% after 48h of treatment **(Figure 5 C)**. The staining of tau (Ser396) decreased subsequent to exposing the cells to Withanolide-A supporting our western blot findings.

### p-Tau Thr 231

At the level of p-Tau Thr 231 hyperphosphorylation, in relation to control again Withanolide-A shows the most significant reduction in the level of hyper-phosphorylation in the site of Thr231 with 40.3% (*p<0.05) and in against of Luteolin and Apigenin the result was not up-to significant with difference of 4.3% (*p<0.05) **(Figure 5 D)**.

## Discussion

Alzheimer’s disease (AD), a progressive neurological illness, is the leading cause of dementia, affecting over 55 million people globally and causing significant financial burden. In terms of therapeutic approaches, there are three main fronts: Approved Drugs-These include aducanumab, donepezil, galantamine, rivastigmine, memantine, and a combination of memantine and donepezil. These drugs temporarily improve symptoms of memory loss and problems with thinking and reasoning. Therapies Under Investigation: These primarily target amyloid-β (Aβ) pathology and tau pathology. They include γ-secretase inhibitors, β-secretase inhibitors, α-secretase modulators, aggregation inhibitors, metal interfering drugs, drugs that enhance Aβ clearance, inhibitors of tau protein hyperphosphorylation, tau protein aggregation inhibitors, and drugs that promote the clearance of tau. Alternative Therapies: These are designed to improve lifestyle, thus contributing to the prevention of the disease [62]. There is still no effective disease-modifying treatment or cure for AD, despite decades of study. The primary goals of the available treatments now are symptom management and quality of life enhancement [63]. Over 99% of candidates in clinical trials for AD treatments fail to receive approval, making the failure rate for these trials infamously high [64]. So, in-silico studies can offer promising solutions to address these challenges and accelerate the development of effective treatments for AD by using computational approaches that provide valuable tools for identifying and validating new therapeutic targets and accelerate the development of safe and effective therapeutic interventions for this debilitating disease, by bridging the gap between experiment and clinical trials, reducing the overall time, cost and number of experiments [65]. The study aims to understand the novel target genes and also their AD-related mechanisms, of *Datura metel* phytochemicals via network pharmacology further the interaction was validated using molecular docking analysis using an in-silico system, with in-vitro study to understand the underlying mechanism and therapeutics approach to the development of new drugs, making the process faster, cheaper and more efficient.

*Datura metel* being toxic in nature researchers have found some potential benefits from different parts of the plants including seeds, leaves, stems, roots, etc. that they can exhibit various pharmacological properties, including antioxidant, anti-inflammatory, and anticancer activities; further studies are required to understand its full potential in humans. Apigenin and Luteolin are flavonoids synthesized through the phenylpropanoid pathway in plants. The pathway involves the conversion of phenylalanine to cinnamic acid, followed by a series of enzymatic reactions leading to the formation of flavones [66] which undergo various metabolic transformations in the human body including glucuronidation, sulfation, and methylation, increasing their water solubility and facilitate excretion [68], whereas withanolide A is a steroidal lactone synthesized from cholesterol through the mevalonate pathway in plants. The pathway involves the conversion of acetyl-CoA to mevalonate and subsequent transformation culminating in Withanolide [67], which under metabolism in humans involves its conversion to various metabolites, including withanolide D, withanolide E, and withanolide G, contributing to various biological effects [69]. Now as the results of the In-silico study suggest, the common pathways that play a role in neuroprotection from Aβ and Tau hyperphosphorylation in Ser 396 and Thr 231 are the IL-17 signaling pathway and TNF signaling pathway which plays a role when *Datura metel* phytochemicals were treated with SH-SY5Y neuroblastoma cells. The result of an In-vitro study in a cell line of SH-SY5Y cells cultured in DMEM media validates the result of Pharmacokinetics studies when the neuroblastoma cells mimic the early-stage pathophysiology of cholinergic neurons affected by AD. One of the studies showed that SH-SY5Y cells differentiated with retinoic acid and BDNF, when challenged with sublethal doses of okadaic acid (OA) and amyloid-beta oligomers (AβOs), provided an in vitro model that mimics the early-stage pathophysiology of cholinergic neurons affected by AD [70]. The differentiated SH-SY5Y cells when treated with Apigenin, Luteolin, and Withanolide-A show overall cell viability in Aβ treated cells were increased in the case of Luteolin and Withanolide-A when the concentration of phytochemical was increased up to 20 (μM), suggesting that these bio-active compound has played a role in one of the above common pathway and inhibit the neuronal death, When the test was repeated with Aβ_1-42_ and p-Tau ser396 & p-Tau Thr231 the affect was decreased by 40% nearly in case of Withanolide A in both p-Tau ser396 & p-Tau Thr231, validating the result of Docking and Network Pharmacology where the highest docking result was shown by Withanolide-A itself suggesting a role in the IL-17 as well as TNF-signaling pathway system for this to act as therapeutic agent to treat Alzheimer’s disease. Now to understand the role and mechanism of how this pathway plays a role in the breakdown of Aβ_1-42_ and p-Tau hyperphosphorylation in ser396 and Thr231, we take the help of various reviews and research articles to underly the actual pathway and mechanism in the treatment of AD. The Interleukin-17 (IL-17) signaling pathway plays a significant role in Alzheimer’s disease (AD). IL-17A, a cytokine, could be crucial in the development of Alzheimer’s disease (AD). Studies have found that AD patients have higher levels of IL-17A in their blood and cerebrospinal fluid (CSF). For instance [71] found increased IL-17A levels in the serum of Chinese patients, while [72] reported increased CSF levels of IL-17A [73] also found that AD patients have significantly higher baseline levels of IL-17A compared to healthy individuals. On a cellular and genetic level, there’s evidence linking IL-17A to AD. Increased differentiation and activation of TH17 cells, along with related transcription factors, have been reported in AD patients [74]. This increase in IL-17A expression could be due to variations in Th17-related genes [75]. BACE1, a protease that contributes to plaque formation in AD, also plays a role. T cells lacking BACE1 have shown reduced IL-17A expression under Th17 conditions in AD mouse models [76]. However, the impact of IL-17A on AD is still a topic of debate. For instance, an AD mouse model with overexpressed IL-17A showed that IL-17A doesn’t worsen neuroinflammation. Overexpression of IL-17A led to a decrease in soluble Aβ levels in the CSF and hippocampus, suggesting our hypothesis that, Withanolide-A is best in Neuroprotection in AD down-regulate this pathway leading to the breakdown of Aβ_1-42_ level in neuronal cells and it also improves glucose metabolism [77]. We further increase our inquisitiveness toward finding the relation between IL-17 and TNF signaling pathways in AD and yes there was, as IL-17 been a pro-inflammatory cytokine that elevates the level of inflammation through the TLR4/NF-κB signaling pathway and microglial activation. Stimulated Microglia express a variety of receptors, including TLRs, which are pattern-recognition receptors [78]. When microglia are stimulated by debris from apoptotic and necrotic cells and heat shock proteins, TLR4 (Toll-like receptor 4) is activated, which further stimulates the downstream transcription factor Nuclear factor (NF)-κB and promotes the secretion of the proinflammatory cytokines TNF-α, IL-1β and IL-6 [79]. So, when IL-17A is increased TNF-α levels in brain is also increased in the brain and exacerbated neuroinflammation through the TLR4/NF-κB signaling pathway indicating that IL-17A may affect the progression of AD by up-regulating the expression of TNF-α in AD brain by regulating the above signaling pathway [80]. As the role of these pathways is highly connected to pro-neuroinflammatory extracellular protein in the membrane, Withanolide-A has shown some very promising activity in decreasing the neuro-inflammation, restoring Glutathione levels, reducing Beta-Amyloid Plaque Aggregation, Regulating Heat shock proteins, and also in inhibiting Oxidative and Inflammatory constituents. Restoring Glutathione levels in the hippocampus that are depleted due to hypoxia, where glutathione is a potent antioxidant that protect cells from damage caused by reactive oxygen species, which are increased in neuro-inflammation [81]. Research that has been published shows that in cultured normal rat cortical neurons, derivatives such Withanolide A increase the expression of α-secretase and decrease the expression of β-secretase. The administration of ADAM10 to APP results in a decrease in Aβ synthesis, since soluble and non-toxic APPα is produced instead. Consequently, in cultured neurons, Withanolide A increased the amount of soluble APPα produced. Additionally, it has been demonstrated that Withanolide A increases the production of an important proteolytic insulin-degrading enzyme (IDE) linked to the breakdown of Aβ. These results suggest that by increasing the synthesis of soluble APPα and Aβ, Withanolide A may reduce Aβ. Withanolide A is a strong contender for a multifunctional Alzheimer’s disease medication given these results. Moreover, withanamide prevented the amyloid plaque-induced mortality of neurons. According to studies using molecular modelling, withanamide A and C bind specifically to the active beta-amyloid pattern (Aβ 25–35) and prevent the development of fibrils [82]

## Materials and Methods

### Experimental

#### Plant material collection

In this research, twenty-five diverse Datura accessions were meticulously studied for their biochemical traits. These accessions were cultivated under controlled environmental conditions using a randomized block design across two crop seasons, 2022 and 2023. To ensure representativeness, five healthy plants per accession were randomly selected and marked for sample collection. Leaves were gathered during the peak vegetative phase, while seeds were harvested at physiological maturity. All samples underwent a controlled drying process for 7-10 days in a sheltered environment with passive air circulation before being preserved in air-sealed containers. Further preparation involved pulverization using a Sujata commercial blender from India and sifting through a Prayag stainless-steel sieve with a 20-mesh size. This meticulous methodology was followed to ensure consistency and accuracy in the subsequent extraction process.

#### Extraction, fractionation, and isolation

After finely grinding the dried leaf samples, 0.5 g was accurately weighed and placed in centrifuge tubes. These specimens underwent extraction using 20 mL of 0.05 M HCl while being agitated for 3 hours at ambient temperature using an orbital shaker. Subsequently, the extracted materials were subjected to centrifugation at 4000 rpm for 10 minutes, leading to the collection of the resultant supernatants. Residues were washed twice with 5 mL 0.05 M HCl, and then centrifuged. The accumulated supernatants were combined and brought to a final volume of 50 mL using 0.05 M HCl within 50 mL volumetric flasks. Before being injected into the HPLC system, the extracts underwent filtration through 0.45 μm syringe filters. The measurement of scopolamine was conducted using an Agilent 1200 series HPLC setup, featuring a C18 column (Zorbax Eclipse XDB, Agilent, 250×4.6mm, 5μm), for accurate quantification. The eluent consisted of a progressive mixture of acetonitrile and a 0.05 M phosphate buffer with a pH of 7.0, flowing at a rate of 1 mL/min. Detection occurred at 215 nm utilizing the diode array detector integrated into the HPLC system, while quantification was accomplished using external standards (scopolamine hydrobromide). Hyoscyamine quantification utilized the Prominence-i HPLC system with a C18 column (Kinetex EVO, Phenomenex, USA). Acetonitrile and phosphate buffer formed the mobile phase. Detection at 215 nm employed the diode array detector, and quantification utilized external standards (hyoscyamine sulfate).

Atropine quantification used an Alliance 2695 HPLC system, with a C18 column (Symmetry Shield, Waters, USA). The composition of the mobile phase involved a mixture of acetonitrile and 0.05M phosphate buffer with a pH of 7. Detection occurred at 215 nm employing the diode array detector, while quantification was dependent on external standard atropine sulfate. Dried seeds had been powdered and exactly weighed samples of 0.1 g were taken into centrifuge tubes. Each sample was extracted with 10 mL of HPLC grade methanol by vertexing for 5 mins using a vortex mixer. The centrifugation of the extracts was carried out at 3000 rpm for 10 minutes.

Subsequently, the resulting supernatants underwent filtration using 0.22 μm syringe filters prior to being injected into the HPLC system. The quantification of Withanolide A was accomplished through Agilent 1200 High-Performance Liquid Chromatography (HPLC), employing a C18 reverse-phase column and a mobile phase of methanol-water. Detection was conducted at a wavelength of 227 nm. Dried seeds were powdered, extracted with methanol as described earlier, and then filtered through Axiva syringe filters before HPLC injection. Quantification employed a C18 column alongside a mobile phase composed of methanol and water, coupled with detection at 227 nm. This process relied on an external standard, specifically Withanolide B. The powdered leaves were extracted with 80% methanol using ultra sonication. Extracts were then centrifuged, filtered, and subjected to UV-Vis spectrophotometry. Absorbance at 272 nm was measured, and quantification utilized external standards: Luteolin and Apigenin. Analysis of variance (ANOVA) employed the R program’s Variability package (Team RC, 2016), treating accessions as fixed effects and environments as random effects. Statistical significance was set at P < 0.001, with result assessment using the least significant difference (LSD) method at p < 0.05 through the R program’s doebio search package. Descriptive statistics including mean, range, and standard error were derived from the same R program package [27].

#### Screening of the metabolites through ADMET Evaluation

The three metabolites that have been isolated, refined, and described must adhere to “Lipinski’s Rule of Five,” which aids in comprehending the physio-chemical characteristics of the substances and determines if it evaluates the intended metabolites’ capacity for absorption in vivo. A compound must meet Lipinski’s Rule of Five in order to be considered a more dependable drug candidate. This rule stipulates that the molecular weight (MW) of the compound should be less than 500, the number of H-bond donors (Hdon) should be less than 5, the number of H-bond acceptors (Hacc) should be less than 10, the (LogP) lipid-water partition coefficient should be less than 5, and the final number of rotatable bonds (Rbon) should be less than 10. The organic metabolites also have to cross the Blood Brain Barrier (BBB) as in the case of cerebo studies and its relatives’ studies [28]. The online tool for swissADMET (Absorption, Distribution, Metabolism, Excretion, and Toxicity) is (http://www.swissadme.ch), which helps us to evaluate the properties of ADME for the desired compounds. LogS, TPSA, and Skin permeation can also be measured through this web tool easily [29]. Prediction of pharmacokinetic and toxicity profiles was carried out online using webform and Protox. It is enough to enter the SMILES code of ligands obtained from PubChem, then click ADMET. Furthermore, the prediction results of these compounds are shown which consist of several parameters of absorption, distribution, metabolism and excretion [30]. Identification of toxicity parameters through the SMART CypP50 server (https://smartcyp.sund.ku.dk/mol_to_som) [31].

#### Collection of Drug Targets

In this study, we employed the Swiss Target Prediction online tool (http://www.swisstargetprediction.ch/) to compile a comprehensive collection of drug targets. Swiss Target Prediction is a widely recognized and reliable resource for predicting potential target proteins for a given set of compounds. To assemble our list of drug targets, we input our selection of compounds into the tool, which then utilized its extensive database and algorithm to predict potential protein targets based on molecular properties and ligand-receptor interactions. This approach allowed us to efficiently identify potential 100 drug targets with high specificity. The resulting list of drug targets forms a critical component of our research, enabling us to explore the mechanisms underlying the therapeutic effects of the compounds under investigation [32]. Now to provide the input search for the Swiss target Prediction all the metabolites were first structured and canonical SMILES for this were obtained from the PubChem database (https://pubchem.ncbi.nlm.nih.gov/) [33].

#### Mining of Genes-Related Targets to AD

To identify specific targets associated with Alzheimer’s Disease, a multifaceted neurodegenerative disorder, we harnessed the power of the Comparative Toxicogenomic Database (CTD https://ctdbase.org), an invaluable online resource that amalgamates the exploration of intricate molecular relationships between chemicals, genes, and diseases through advance understanding about how environmental exposures affects human health. The default parameter for CTD was set to interpret the significance of Alzheimer’s disease by going to disease section, by default, the tool used a corrected p-value threshold of 0.01, and the result was exported in .csv file, for further analysis [34]. DisGeNET (https://www.disgenet.org) for mining and extracting Genetics and Disease-related information, through an extensive array of scientific literature and database The default range for the DisGeNET GDA score is 0.1 to 1. This means that only associations with a GDA score of 0.1 or higher will be considered significant. However, users can also specify a different range for the GDA score, depending on their specific needs [35], GenBank (https://www.ncbi.nlm.nih.gov/genbank/) identify a multiple of genes and genetic sequences with implications in AD pathogenesis The GenBank Genome Data Assembly (GDA) score is a quality metric assigned to genome assemblies submitted to GenBank. It ranges from 0 to 100, with higher scores indicating higher quality assemblies [36], and last we utilized the Online Mendelian Inheritance in Man (OMIM) (https://www.omim.org) online tool, a reputable resource that compiles a wealth of genetics and clinical information The default GDA score range is 0.05 to 1.0. This means that genes with a GDA score of 0.05 or higher are considered to be potentially disease-associated. Genes with a GDA score below 0.05 are considered to be unlikely to be associated with a human genetic disease [37]. By searching for relevant keywords, such as ‘Alzheimer’s Disease,’ ‘Neurodegenerative disorders, ‘and’ AD-associated Proteins, ‘along with the screening only based on “*Homo sapiens*”.

#### Intersection of Gene Targets

To obtain these we used a web-based online tool for the intersection between gene targets of Datura and Alzheimer’s disease using Venny 2.1 (https://bioinfogp.cnb.csic.es/tools/venny/index.html), the common intersected targets were further analysed to obtain possible targets for AD treatment [38].

#### Protein-Protein Interaction (PPI) Enrichment and Clustering Analysis

Using the STRING database version 11.0 (https://string-db.org/), a PPI network was created. Only the minimum needed interaction score > 0.4 was selected as significant, the Network type was selected as Full Network, and with maximum additional interactor was kept at just zero and the organism set to *Homo sapiens*. PPI networks are made up of nodes, which stand in for a target protein, and edges, which stand in for interactions between proteins. The total score is represented by an edge’s thickness. The degree of a node describes how many other nodes are directly related to it. The importance of a node increases with degree [39]. Using Cytoscape software (v.3.7.1) and its Network Analysis plugin, core targets were discovered using network analysis. The top 10 proteins, ordered by degree, were chosen and designated as key targets in the current investigation [40].

The PPI network was analysed for clustering modules using the MCODE plugin. The selection criteria for MCODE included a degree cut-off of 2, node score cut-off of 0.2, k-core of 2, and max depth of 100. MCODE identified the node with the highest weighted vertex as the seed node, which was the main target of the cluster [41]. Furthermore, a network for potential alkaloid target-AD target was created using Cytoscape software [40].

#### GO Function and KEGG Pathway Enrichment Analysis

Metascape (https://metascape.org/gp) is a comprehensive tool used for gene annotation and enrichment analysis. The analysis of GO biological process and KEGG pathway enrichment was conducted using Metascape. Significant enrichment words had a p-value of less than 0.01, a minimum count of 3, and an enrichment factor of more than 1.5. The top 10 enriched terms were visualized using an online tool [42]. Additionally, a KEGG pathway network for the main metabolites against AD was created using Cytoscape software. Red nodes represent enriched KEGG pathways, while brown nodes represent target proteins [43].

#### Determination of Critical Potential Targets Based on CytoHubba-Alzdata Database

The data on drugs, components, targets, pathways, and diseases was entered into Cytoscape 3.9.0 for key component and target analysis. Maximal Clique Centrality (MCC), Degree, and other network properties are examined using the CytoHubba plug-in’s features where the top Hubba nodes parameter were chosen as Top 10 and the display option was set to Display the shortest path, considering the goals that satisfy the MCC and degree requirements as potentially crucial goals [44]. AlzData (https://www.alzdata.org). An AD database that gathers recent high-throughput omics data. The AlzData database was used to analyse correlations between the pathophysiology of AD (Aβ and tau) and the gene symbols for the target proteins of UR alkaloids against AD. The collected results were then organized using Microsoft Excel. We conducted further GO and KEGG pathway enrichment studies using the targets of the UR alkaloids associated with AD pathogenesis. In the control and AD groups of the GEO dataset, the “Differential expression” Entorhinal cortex, Hippocampus, Temporal cortex, and frontal cortex module of AlzData were examined the normalized expression targets of phytochemicals against AD [45]. Higher scores and darker colorations showed a more significant link between the genes and AD. The MCC algorithm was used to produce scores representing the strength of relationships between nodes and edges. The top 10 target genes/proteins with the highest scores for each active ingredient were then identified. Additionally, the Panther categorization system (http://pantherdb.org/) was used to classify the target proteins. The core Protein-Protein Interaction network cluster was then identified by using the Cytoscape plugin MCODE to analyse the relevant network [46].

#### Molecular Docking

For the purpose of studying the interactions between receptor and ligand, docking was used to control the molecular docking performance. The PDB database (http://www.rcsb.org) was used to get the protein structures of hub genes [47], and PubChem (https://pubchem.ncbi.nlm.nih.gov/) was used to obtain the mol2 file formats for the molecular structures of Selected metabolites [33].

PyMOL software was utilized to eliminate excess inactive ligands and water molecules [48]. The proteins were then charged and hydrogenated in AutoDockTools1.5.6 software before being exported to pdbqt format [49]. AutoDock Vina software was used to conduct molecular docking simulation of potential targets and their corresponding components, following established methods [50]. Docking results were visualized using LigPlot+ (http://www.ebi.ac.uk/thornton-srv/software/LIGPLOT/) and PyMol (Version 4.6.0), and the interactions were further visualized using LigPlot+ [51].

#### Molecular Dynamics (MD) Simulation

Following the initial evaluation of binding affinity and hydrogen bonds, the selected drug-receptor complexes underwent Molecular Dynamics Studies (MDS) employing GROMACS. The Molecular Dynamics (MD) simulation utilized GROMACS v2016.16 and the CHARMM 27 force field [52], focusing on three protein structures: plasmin-textilinin-1 (PDB: 3UIR), Novel c-Jun N-Terminal Kinase (JNK) (PDB: 2P33), and Cytochrome P450 (PDB: 3EQM) in conjunction with Withanolide-A. Ligand topologies were obtained from SwissParam (https://www.swissparam.ch/), and simulations adhered to a standardized protocol [53]. This protocol involved the creation of a triclinic water box, solvation with ’spc216-Simple Point Charge water,’ introduction of NaCl at a concentration of 150 mM for system neutralization, and subsequent system minimization [54].

The system underwent energy minimization, followed by equilibration in the NVT ensemble (constant number of molecules, volume, and temperature) for 0.1 ns. Subsequently, a 100 ns simulation was conducted in the NPT ensemble at 300 K [55]. Protein-ligand coupling underwent a two-step equilibration, NVT and NPT, each lasting 10 ns, utilizing the V-rescale modified Berendsen thermostat. The LINCS algorithm was applied to constrain the lengths of hydrogen-containing bonds [56]. 100 ns Molecular Dynamics (MD) run with GPU acceleration using CUDA Toolkit 12.2 (https://developer.nvidia.com/cuda-downloads) was executed under periodic boundary conditions. The analysis of results employed VMD and Chimera, focusing on parameters such as RMSD, RMSF, Rg, interaction energy, hydrogen bonds, and their distances. Graphical representations were generated using GRACE [57].

#### Cell Culture and Treatments

Human SH-SY5Y neuroblastoma cells were cultured in Dulbecco’s Modified Eagle Medium, supplemented with 10% fetal bovine serum and 1% penicillin-streptomycin at 37° C in 5% CO2. For assays, cells were seeded in 96-well plates at a density of 1Χ10^4^ cells/well and allowed to adhere overnight [58]. The following compounds were tested: luteolin, Apigenin, and Withanolide A (all purchased from Sigma-Aldrich, USA). Compounds were dissolved in DMSO and added at concentrations of 5μM, 10μM, and 20μM for 24 hours. Amyloid-beta (AnaSpec, USA) was added at 10μM for 24 hours to induce toxicity. Equal volumes of DMSO served as vehicle controls.

#### Cell viability assay

Cell viability was assessed by the MTT (3-(4,5-dimethylthiazol-2-yl)-2,5-diphenyltetrazolium bromide) reduction assay. MTT was added to cells at a final concentration of 0.5mg/mL and incubated for 3 hours. Formazan crystals formed were dissolved in DMSO and absorbance was read at 570nm using a microplate reader (BioRad, USA). Viability was calculated as a percentage of DMSO vehicle control [59].

#### Amyloid-Beta 1-42 ELISA Test

The conditioned medium was collected after treatments and centrifuged at 2000xg for 10 minutes. Levels of secreted amyloid-beta 1-42 were quantified using a sandwich ELISA kit (Invitrogen, USA) as per the manufacturer’s protocol. Absorbance was measured at 450nm [60].

#### Western Blot for Tau Phosphorylation

After treatments, cell lysates were prepared using RIPA buffer containing protease and phosphatase inhibitors. Protein concentrations were estimated by BCA assay (Thermo Scientific, USA). Equal amounts of proteins (30μg) were resolved on 10% SDS-PAGE gels and transferred to PVDF membranes. Membranes were blocked with 5% non-fat milk and incubated overnight with primary antibodies for p-Tau Ser396, p-Tau Thr231, and beta-actin (Cell Signaling Technology, USA) followed by HRP-conjugated secondary antibody. Bands were visualized using ECL substrate (BioRad, USA) and analysed by densitometry [61].

## Data Availability Statement

Data associated with this study are not available, as this study does not require any ethical clearance.

## Acknowledgement

We would like to express our sincere gratitude to Parul University, Ranchi University and Chaudhary Charan Singh Haryana Agricultural University for their invaluable contributions to this research project. Their support, guidance, and expertise have been instrumental in the completion of this work.

## Author contributions

In this study, [MS] primarily led the conceptualization and [MJ], who developed the initial idea and research framework. The methodology design was a collaborative effort between [MS & MJ] and [MS,MJ,AKD,PK], ensuring the study’s methodological rigor. Data collection responsibilities were undertaken by [NP] and [AB,DT], who meticulously gathered the necessary information for analysis. The data analysis phase was spearheaded by [NP] and [AB], who applied statistical methods to interpret the findings. Writing the original draft was a joint effort involving [MJ], [MS], and [AKD,PK], while all authors contributed to reviewing and editing the manuscript. Visualizations were crafted by [NP], adding a visual dimension to the research outcomes. Supervision of the entire research process was overseen by [MJ & PK], ensuring quality control and adherence to research standards. Lastly, funding acquisition was successfully managed by [MJ & PK], securing the financial support necessary for the study’s completion.

## Declaration of interest statement

There is not conflict of interest among the authors.

## Notes

### Competing Interest Statement

The authors have declared no competing interest.

